# Acto-myosin cross-bridge stiffness depends on the nucleotide state of the myosin II

**DOI:** 10.1101/2020.07.08.191593

**Authors:** Tianbang Wang, Bernhard Brenner, Arnab Nayak, Mamta Amrute-Nayak

## Abstract

How various myosin isoforms fulfill the diverse physiological requirements of distinct muscle types remains unclear. Myosin II isoforms expressed in skeletal muscles determines the mechanical performance of the specific muscles as fast movers, or slow movers but efficient force holders. Here, we employed a single-molecule optical trapping method and compared the chemo-mechanical properties of slow and fast muscle myosin II isoforms. Stiffness of the myosin motor is key to its force-generating ability during muscle contraction. We found that acto-myosin (AM) cross-bridge stiffness depends on its nucleotide state as the myosin progress through the ATPase cycle. The strong actin bound ‘AM.ADP’ state exhibited > 2 fold lower stiffness than ‘AM rigor’ state. The two myosin isoforms displayed similar ‘rigor’ stiffness. We conclude that the time-averaged stiffness of the slow myosin is lower due to prolonged duration of the AM.ADP state, which determines the force-generating potential and contraction speed of the muscle, elucidating the basis for functional diversity among myosins.

## Introduction

Skeletal muscles account for roughly 40 % of the human body weight, and undertake several physiological roles, including movement and maintaining the body posture. Different skeletal muscles exhibit a wide range of muscle performance. For example, muscle responsible for bearing the load or maintaining the posture display slower shortening velocity as compared to the muscles dedicated for rapid movements. The diverse mechanical performance of muscles is attributed to various isoforms of the main force generating motor proteins, myosin II. It is believed that the varied force transduction capacity of the myosin heavy chain (MHC) isoforms expressed in distinct muscle types determines their specialized physiological role. During muscle contraction, myosin exploits the energy derived from ATP hydrolysis to drive mechanical work. ATP driven cyclical interaction between the myosin and actin filaments generate force and causes muscle shortening. Acto-myosin (AM) ATPase cycle proposed from biochemical studies (White and Taylor, 1976) comprises different nucleotide states of the myosin active site, exhibiting strong and weak interactions with actin. Accordingly, the mechanical interaction between actin and myosin begins with ADP.Pi state; subsequent Pi release is associated with the generation of powerstroke and formation of AM.ADP strong bound state, ADP release from myosin active site results in an AM rigor state. The rigor state ends when ATP binds to the myosin causing acto-myosin to detach.

The class II skeletal myosin isoform-MHC-1 is primarily expressed in slow skeletal, aerobic muscles, while MHC-2 is found in white, anaerobic, fast contracting muscle. Slow muscles develop lower force, slower unloaded shortening velocity, and a comparable or higher thermodynamic efficiency than the fast muscle, corresponding to its physiological role to maintain isometric force with relatively low energy consumption (Barclay et al., 1993, He et al., 2000). The MHC-1 and MHC-2 are endowed with unique sets of essential (ELC) and regulatory lights chains (RLC) associated with the heavy chains, which provide structural stability to the lever arm and fine-tunes the motor activity (Lowey and Trybus, 1995, Nayak et al., 2020).

Almost 10 fold difference in the mechanical characteristics, such as unloaded shortening velocity of the fast and slow myosin isoforms have led to several investigations including, studies of kinetics and mechanical properties as well as structure-function analysis. The isoform-dependent functional variations are being extensively studied by measuring the kinetic differences in the fast and slow myosin isoforms for several species including rat, rabbit, and human. Despite following the same kinetic cycle, the two isoforms were shown to primarily differ in their acto-myosin detachment rates, which are limited by the rate of ATP binding for the fast myosin and by the rate of ADP release for slow myosin. From kinetics studies, difference in the ADP release rate was concluded to be a major factor responsible for the differences in the shortening velocity, and slow ADP release linked to slower velocities for the MHC-1 isoform (Siemankowski and White, 1984, Iorga et al., 2007).

Apart from the kinetic properties, a major difference in the mechanical properties reported to be the force-generating capacity of the two isoforms, which was found more than two fold different. The resistance to elastic deformation, i.e., stiffness of the force-generating acto-myosin cross-bridges was proposed to be central for efficient force production and displacement (Huxley and Simmons, 1971, Eisenberg and Hill, 1978). Besides, as the molecular stiffness modulates the deformation in the head domain by loads acting on the lever arm, stiffness is expected to couple catalytic activity with the load. Single molecule studies of rat myosin isoforms provided precise details into kinetic and mechanical properties, including the difference in the acto-myosin detachment rates and the elasticity (Capitanio et al., 2006). In this study, the slow myosin isoform was reported as more compliant than the fast myosin isoform. Generally, the single-headed myosin S1 (Subfragment-1) employed in single-molecule investigation is a preferred -minimal, but sufficient motor component. One concern however is that the proteolytic enzymes, papain or chymotrypsin employed to generate single-headed myosin S1 cause a partial loss of either ELC or RLC, respectively.

In accordance with single-molecule studies, single muscle fiber analyses found at least 2-3 fold lower isometric force developed by slow muscle fibers than by fast ones. Yet again, lower stiffness for slow myosin was estimated than the fast isoform. In these studies, equal number of active acto-myosin cross-bridges in slow and fast fibers during isometric contraction was reported (Brenner et al., 2012), reinforcing the notion that the individual motor head stiffness varied, and consequently the force generation by the two myosin isoforms. Along similar lines, about 60% more myosin was shown to engage in slow muscle fibers (Percario et al., 2018) to attain same amount of force as the fast fibers. Both the single molecule and single muscle fiber studies estimated the stiffness of ~0.5 pN/nm, and 1-2 pN/nm for slow MHC-1 and fast MHC-2 myosin molecules, respectively (Percario et al., 2018, Brenner et al., 2012, Capitanio et al., 2006). Thus, in addition to the kinetic difference, stiffness of the myosin molecules is a highly discussed parameter and proposed to be a major determinant for the difference in the mechanical performance of various muscle types (Brenner et al., 2012, Percario et al., 2018).

Despite the well-designed studies in muscle fibers and at the single-molecule level, open questions remain, i.e., 1) whether single myosin heads employed in single-molecule studies recapitulate the characteristics of full-length dimeric myosin motors, and 2) if the elasticity of single molecules, estimated from the cohort in the sarcomeres in single fibers studies is a true representation of the single myosin property.

There are no studies involving the direct comparison of rabbit fast and slow skeletal myosins in the single-molecule investigation methods. Here, we employed optical trapping of individual native full-length motor molecules to examine the isoform-specific chemo-mechanical properties of the rabbit MHC-1 and MHC-2. We characterized the acto-myosin detachment kinetic rates and found that MHC-1 exhibits more than 15 fold slower ADP release rate compared to the MHC-2. The two isoforms displayed significantly different powerstroke size with MHC-1 supporting longer mean displacement of nearly 6 nm. Intriguingly, in contrast to previous studies, we have observed similar ‘rigor’ stiffness for both the myosin isoforms. Detailed analysis of MHC-1 led us to identify two distinct AM cross-bridge stiffness. We found nucleotide states dependent cross-bridge stiffness, i.e., MHC-1 in AM.ADP ‘strongly bound’ and AM ‘rigor’ state displayed ~2 fold difference. Apart from the dimeric motors, the single headed myosin also exhibited this AM cross-bridge state dependent difference in the elasticity. Our results suggested that the duration of the strong bound ADP state defines the measured stiffness for the MHC-1 and therefore the rate of ADP release remains the main factor defining shortening velocities.

Our findings provide new insights into understanding the fundamental mechanism of muscle type -specific force-generating potential and contraction speed.

## Results

### Fast *vs* slow myosin isoforms, ATPase kinetics

We isolated myosin II motors from rabbit fast-twitch, *Musculus psoas,* and slow-twitch, *Musculus soleus* muscle fibers to investigate the two myosin isoforms that leads to a diverse mechanical properties i.e., force generation and movement. Rabbit *psoas* muscle is known to expresses mainly the fast isoforms MHC-2×/2d (~92%) and MHC-2b (~8%), whereas *soleus* muscle contains almost exclusively MHC-1 (~97%) (Tikunov et al., 2001). We inspected the native fast and slow myosin extracted from respective muscles in the SDS-PAGE gels to ensure the purity (Figure S1). MHC-2 is henceforth referred as psoas fast myosin (^Pso^M-II) and MHC-1 referred as soleus slow myosin (^Sol^M-II). Using *in vitro* actin filament gliding assay, we assessed the velocity of actin filaments driven by the two myosin isoforms. The two motors exhibited >10 fold difference in the actin filament gliding speed driven by the respective motors (cf Figure S1) i.e., 3.73 ± 0.047 μm/s *vs.* 0.265 ± 006 μm/s, under identical experimental conditions (2 mM ATP, 22°C). This result is consistent with the difference observed in the shortening velocity of fast *vs.* slow skeletal muscle fibers (Larsson and Moss, 1993).

The shortening velocity in muscle fibers or actin filament gliding velocity driven by isolated motors are primarily determined by the duration of strongly bound acto-myosin states ‘t_on_’ and the stroke size (δ) of the myosin during acto-myosin cross-bridge cycle.

We probed the differences in the two myosin isoforms for these velocity determining parameters using single molecule analysis approach. With the optical trapping method introduced by Finer et al (Finer et al., 1994), it is possible to measure the kinetic and mechanical parameters of the individual motor molecules to understand even the subtle differences between the motor properties that may ultimately influence the ensemble motor behavior. The single molecule analysis approach allowed kinetics and mechanical characterization of motors with a high temporal and spatial resolution.

The optical trapping experimental set up as described previously (Steffen and Sleep, 2004, Steffen, 2006), is schematized in figure 1A. An actin filament suspended and held taut between the two optically trapped beads interacted with the myosin immobilized on the nitrocellulose coated glass bead. The individual bead positions were recorded using quadrant detectors to monitor periodic acto-myosin interactions. As shown in Figure 1B, the well isolated intermittent interactions between the actin and myosin are indicated by the decrease in the Brownian noise of the actin filament dumbbell as the myosin proceeds through the ATPase cycle. As the interaction of the myosin with actin is associated with the generation of a powerstroke, the duration of the reduced noise represents the post-powerstroke strong bound ‘AM.ADP’ and ‘AM’ rigor states. This description of the bound states is valid if the Pi is released before or during the powerstroke. Recent studies have indicated that the powerstroke precedes the Pi release step from the active site (Muretta et al., 2015, Woody et al., 2019). However, duration of AM.ADP.Pi state should be sufficiently long after the powerstroke to consider it as a ‘strong bound state’ contributing to the lifetime of the attached state - (at least 2 ms i.e., the detection limit with our current experimental set up). We cannot distinguish between strong bound AM.ADP.Pi and AM.ADP states. For these reasons, we consider that the AM bound state primarily includes the duration of ‘ADP bound’ and ‘rigor’ state.

**Figure 1.**
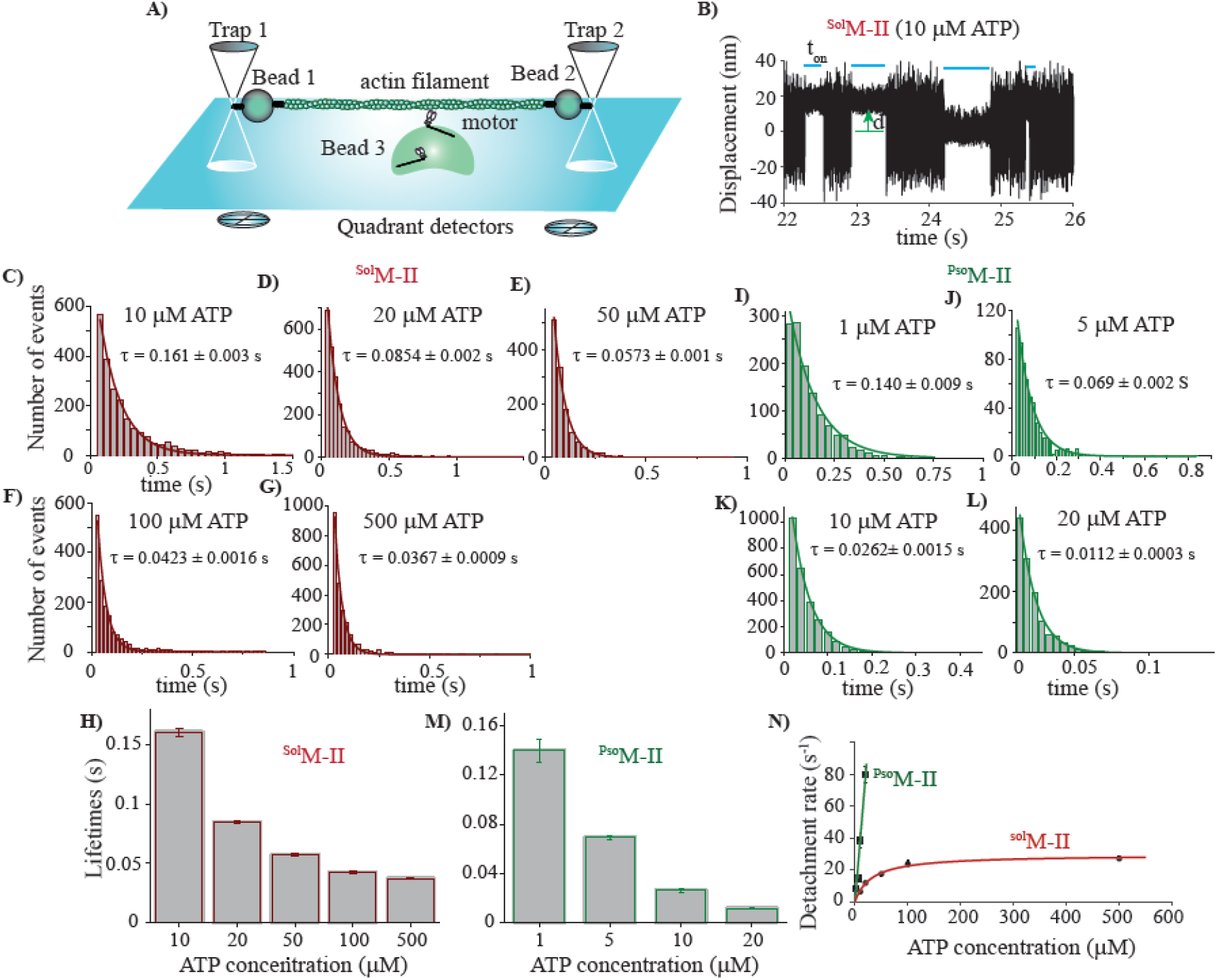
ATP concentration dependence of AM bound lifetimes for ^Pso^M-II and ^Sol^M-II. **A)** Optical trap set up for 3 bead assay. The springs illustrate the elastic elements. The motor immobilized on the 3^rd^ bead is shown in the schematic. Note that the various components of the set up are not drawn to scale. **B)** Original data trace displaying the bead position signal over time. The AM interaction events are indicated with blue horizontal lines. The green arrow illustrates the displacement (d) from average unbound to bound position. **C-G)** Histograms of AM bound lifetimes measured at increasing concentration of Mg-ATP from 10 μM to 500 μM for ^Sol^M-II. The average lifetimes (τ) were calculated by least square fitting each histogram with single exponential decay function. **H)** Bar diagram comparing τ from the fits at different ATP concentrations. Error bars – standard error of the fits. **I-L)** For ^Pso^M-II, the AM bound lifetimes measured from 1-20 μM ATP. Each histogram was fitted with single exponential component to yield average time constant. **M)** Bar diagram displaying τ at various ATP concentrations for ^Pso^M-II. **N)** AM detachment rate (1/τ over different ATP concentration for ^Pso^M-II and ^Sol^M-II, fitted with linear regression and Michaelis–Menten function (r=0.99), respectively. For ^Pso^M-II, k_T_ from linear regression - 3.9634 ± 0.308 μM^−1^ s^−1^, For ^Sol^M-II, kcat = 29.39 s^−1^, Km = 31.58 μM. Altogether, 115 individual ^Sol^M-II molecules, and ~15,000 AM binding events were identified and analyzed. At 10 μM ATP, N = 22, n= 2637; at 20 μM ATP, N = 20, n =3132; at 50 μM ATP, N = 16, n =2356; at 100 μM ATP, N = 23, n =3062 and at 500 μM ATP, N = 34, n =3458. The experiments were performed with myosin molecules from at least three independent ^Sol^M-II preparations. For ^Pso^M-II, 30 individual myosin molecules were measured. At 1 μM ATP, N = 4, n = 1223; 5 μM ATP, N = 6, n = 604; 10 μM ATP, N = 10, n = 2675 and 20 μM ATP, N = 10, n =2296. The experiments were performed with two independent preparations of ^Pso^M-II. N = number of individual myosin molecules and n = number of AM association events. The nonparametric Mann-Whitney U test yielded the statistical differences in t_on_ with P<0.0001 when the event lifetimes for ^Sol^M-II and ^Pso^M-II were compared between different ATP concentrations.

Varying concentration of ATP in these experiments can be used to measure the kinetics of cross-bridge cycling i.e., the duration of AM bound state (t_on_) or the detachment rate (1/ t_on_). Increasing ATP concentration decreases the lifetime of the AM rigor state. Therefore, at saturating ATP concentration ADP release rate can be estimated as this state becomes rate limiting. We modulated the duration of the rigor state within the acto-myosin bound lifetime for slow myosin, ^Sol^M-II, by increasing ATP concentrations. For the fast myosin however, only the rigor state is captured as the ADP release is too fast to be detected with current temporal resolution of our set up (~1 ms).

For ^Sol^M-II, the duration of strong bound states was measured at the range of ATP concentrations from 10 μM - 500 μM. As shown in Figure 1C-H, t_on_ decreased with increasing ATP concentration. The reciprocal of the average t_on_ yields the detachment rate of acto-myosin at specified ATP concentrations as shown in Figure 1N. ATP concentration dependence of the detachment rates for the ^Sol^M-II followed a hyperbolic curve, suggesting attainment of a rate-limiting ADP release step (Figure 1N). For ^Sol^M-II, the hyperbolic fit to detachment rates as a function of ATP concentration yielded maximum AM dissociation rate (k_cat_) of 29.39 s^−1^, and Km of 31.58 μM, that is a measure of ATP affinity. The ADP release rate estimated from the lifetimes at 500 μM ATP concentration was 27.7 s^−1^.

For ^Pso^M-II, the t_on_ was measured at 1 μM - 20 μM ATP concentrations (Figure I-M). Measurements above 20 μM ATP concentrations were not reliable as the shorter events (< 2 ms) became undetectable and very few binding events were observed. The linear increase of detachment rates indicated ATP binding as rate-limiting in this range of ATP concentration (Figure 1N). From linear regression, second order rate constant for ATP binding, K_T_, was estimated to be 3.96 ± 0.3 μM^−1^s^−1^. The K_T_ is similar to the value found in previous measurement for single full length myosin molecules, and for acto-S1 ATPase in solution studies (White and Taylor, 1976) (Takagi et al., 2006).

Taken together, more than 10 fold slower AM detachment rate, indicative of slow ADP release was observed for ^Sol^M-II than ^Pso^M-II. This result was consistent with our previous solution kinetics studies for single headed myosin subfragment 1 (S1) (Nayak et al., 2020).

Collectively, ^Sol^M-II exhibited more than 15 fold slower ADP release rate compared to the ^Pso^M-II. We previously determined the ADP release rate of > 500 s^−1^ for single headed fast myosin (Nayak et al., 2020).

### Fast *vs* slow myosin isoforms, mechanics

As an important velocity determining factor, we further investigated the acto-myosin interactions for force generating conformational change causing actin filament displacement i.e., the powerstroke size (δ) for the two myosin isoforms. Using the ‘shift of histogram’ method, introduced by Molloy et. al. (Molloy et al., 1995), we determined the average stroke size at different ATP concentrations. To our surprise, we found that the measured average stroke size decreased with increasing ATP concentrations (Figure. 2 A-F), for ^Sol^M-II, about 6 nm at 10 and 20 μM ATP decreased to 4.37 nm at 500 μM ATP. For ^Pso^M-II, the stroke size was about 4.5 nm at 1, 5, and 10 μM ATP (figure 2 G and H). Thus, we found statistically significant difference in the mean displacement of fast and slow motors, even for the native two-headed myosins, but only at low ATP concentrations (10 μM ATP). Thus, consistent with a previous report (Capitanio et al., 2006) for rat fast and slow skeletal myosin II-S1, the average working stroke for ^Pso^M-II was smaller than for ^Sol^M-II at specified ATP concentration.

**Figure 2.**
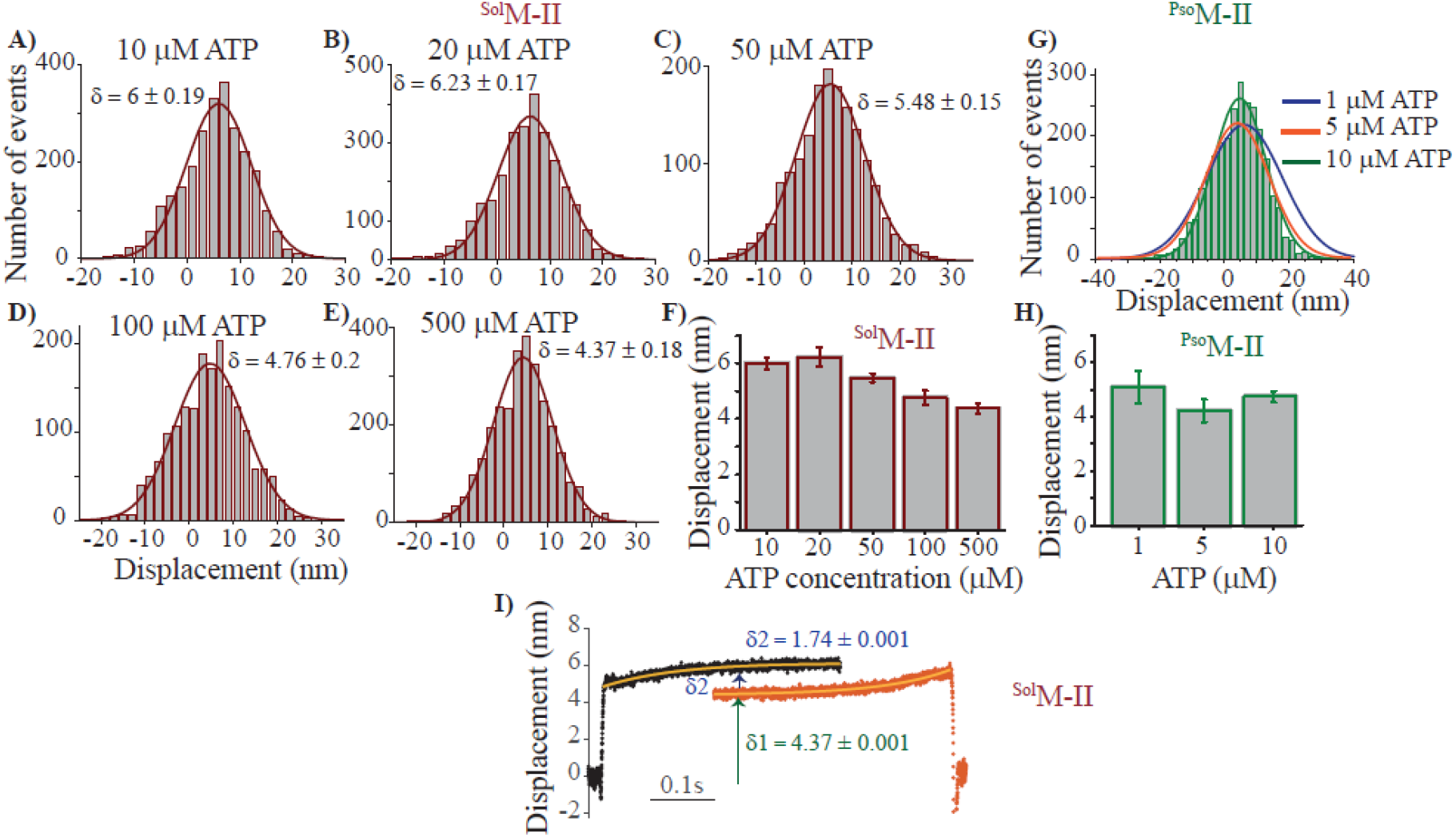
Powerstroke size of ^Pso^M-II and ^Sol^M-II. The average stroke size/ mean displacement was determined by histogram shift (δ) from mean free dumbbell noise. **A-E**) Histograms of displacements measured at increasing concentration of ATP from 10 μM to 500 μM for ^Sol^M-II. The average stroke size (δ) was calculated by least square fitting each histogram with Gaussian function. **F)** Bar diagram comparing the average stroke size at different ATP concentrations. Error bars - standard error of the fits. **G and H)** Stroke size of ^Pso^M-II at 1, 5 and 10 μM ATP. The difference in the power stroke size was statistically significant for ^Sol^M-II between ATP concentrations 10 and 500 μM (P < 0.0001, two-sample independent t-test). Whereas, for ^Pso^M-II the power stroke sizes at ATP concentrations 1 and 10 μM showed no statistically significant difference (P = 0.6, two-sample independent t-test). **I)** Ensemble-average of binding events for ^Sol^M-II, acto-myosin attachment events measured at 10 μM ATP concentrations. The beginning (black data points) and end (orange data points) of the 500 events were synchronized. The beginning and end of the event traces were fitted with single exponential function. ^Sol^M-II showed two steps in the bound events, i.e., δ1 =4.37 ± 0.001 nm and δ2 = 1.74 ± 0.001 nm, respectively. δ2 was calculated by subtracting 1^st^ step from the overall estimated displacement of 6.11 ± 0.002 nm. Important to note that for ensemble averaging, all the AM interactions events were adjusted so as to have same duration (0.4 s). For the analysis, attachment events with a lifetime of at least 0.05 s or longer were selected. The shorter events were prolonged to 0.4 s, and the noise trace of 0.02 s from both the ends (initial AM binding and release) were used. The method is described in detail in Veigel et al.(Veigel et al., 2002).

The possible explanation for the two different stroke size values for ^Sol^M-II, is that at low ATP concentration the AM rigor state dominates ‘t_on_’, thereby the amplitude of displacement measured at low ATP concentrations represents the overall size of the working stroke, whereas at higher ATP concentration the lifetime of the AM rigor state is too short, and AM.ADP state dominates, thereby the displacement comprises primarily the mechanical event associated with Pi release.

Thus, we were able to identify the force-generating power strokes accompanying the hydrolysis product release, i.e., Pi release resulting into large stroke (~ 4 nm) and the ADP dissociation driving the second smaller stroke (~ 2 nm) (Figure 2F). For ^Pso^M-II, however the complete mechanical event was observed, resulting into overall average displacements between 4.27 – 5.08 nm at 1 to 10 μM ATP (Figure 2G and H).

We further intended to resolve the sub-steps within individual bound states. We employed ensemble averaging approach to identify the conformational changes associated with Pi and ADP release, as previously used for different myosin isoforms (Veigel et al., 2003, Capitanio et al., 2006, Greenberg et al., 2014). For ^Sol^M-II, we selected the original acto-myosin interaction records at 10 μM ATP where the event durations were sufficiently long (average duration of 161 ms as shown in Figure 1C). As indicated from our kinetics analysis, bound duration contained both AM.ADP and AM rigor states at this ATP concentration. ‘t_on_’ from about 500 acto-myosin attachment events was aligned at the start and end of event (Figure 2I). For each event, the start and end were precisely determined by thresholding the variance of the original data trace. With this approach the mechanical events of ^Sol^M-II could be resolved into two distinct steps. Amplitudes of the working stroke at the beginning and the end was determined by exponential fitting of the data, yielding the substeps of 4.37 ± 0.001 nm and 1.74 ± 0.002 nm. The stroke size for the respective substeps estimated with this approach supported our results obtained at low and high ATP concentrations. For ^Pso^M-II, however such analysis was not feasible as the ADP bound duration of the acto-myosin was too short even at 1μM ATP, and higher temporal resolution would be needed to assemble enough data points necessary for such an approach.

### Stiffness of fast *vs* slow myosin isoforms

The stiffness of the motor molecules directly influences the generation of force, and is crucial to understand the mechanisms underlying the force generation capacity of respective muscle type. Apart from about 80% amino acid sequence homology in the two MHC isoforms, unique pairs of light chains i.e., essential light chain MLC1f/MLC3f and regulatory light chain MLC2f, associate with ^Pso^M-II, whereas ^Sol^M-II assembles with ELC, MLC1sa /MLC1sb and RLC, MLC2v/s. It is assumed that variants of light chains provide different stiffness to the myosin head. Elasticity of myosin head is expected to affect the mechanical strain of the acto-myosin cross-bridge. The change of strain under load affects the kinetics of some steps of the ATPase cross-bridge cycle (Huxley and Simmons, 1971, Reconditi et al., 2004). For myosin V and smooth muscle myosin, the strain dependence of ATPase kinetics is well reported (Veigel et al., 2003, Veigel et al., 2005).

To determine the stiffness of the ^Pso^M-II and ^Sol^M-II motors, we recorded AM interaction by applying positive position-feedback on the laser-trapped beads as described in Steffen et al (Steffen et al., 2001, Steffen, 2006). Using analog positive feedback in AC mode, the noise amplitude of the free dumbbell was increased. As a consequence, the variance ratio between binding events and free dumbbell noise improved for both traps in the direction of the actin filament axis (x-axis), particularly important for protein with lower stiffness. This procedure allowed event detections and stiffness measurements by increasing the amplitude of Brownian noise by ~30%, without affecting the trap stiffness in the y and z directions.

Using bead variance-covariance method as described previously (Lewalle et al., 2008, Smith et al., 2001), the stiffness of individual motors was calculated as shown in Figure 3. For ^Sol^M-II myosin, we analyzed AM interaction records obtained at various ATP concentrations of 10 μM - 500 μM ATP (Figure 3C). As shown in the scatter plot, stiffness measured at 10 and 20 μM ATP showed large variations, ranging from 0.8 to 2.6 pN/nm. To our surprise, we observed that depending on the ATP concentration the estimated average stiffness of the myosin head was >2 fold different i.e., 1.5 ± 0.11 pN/nm *vs* 0.51 ± 0.05 pN/nm at 10 μM and 500 μM ATP, respectively. As the average duration of AM bound state are about 7 times longer for ^Sol^M-II at 10 μM ATP, we could rule out the presence of contaminant fast myosin in the measurements. The ^Pso^M-II displayed an average stiffness of 1.26 ± 0.17 pN/nm, with no significant difference in the stiffness at 1 and 10 μM ATP (Figure 3D). In fact, the stiffness values were comparable for ^Sol^M-II and ^Pso^M-II at 10 μM ATP, and the disparity was observed only at high ATP concentration for ^Sol^M-II.

**Figure 3.**
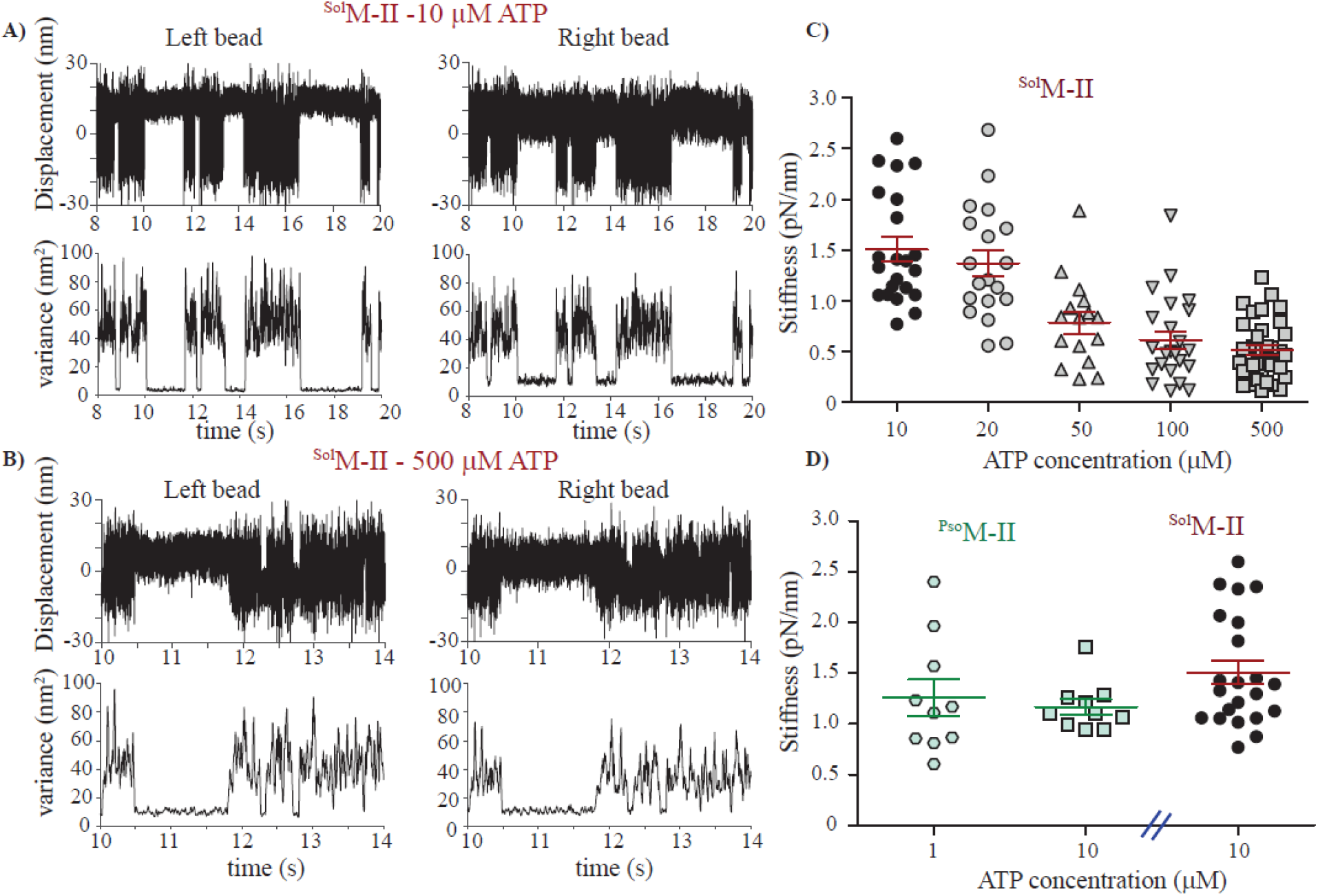
Stiffness of ^Pso^M-II and ^Sol^M-II measured at various ATP concentrations. **A and B)** Original data records acquired in optical trapping experiments for ^Sol^M-II at low (10 μM) and high (500 μM) ATP concentrations. Top traces in each panel shows bead displacement (nm) plotted over time for both left and right bead of the dumbbell. The data record displays 6 binding events **(A)**. Lower traces display corresponding variance of bead displacement as a function of time. The difference in the variance from 2 beads as well as among two different ATP concentrations is apparent. Typically, lower variance ratio as in **(B)** was observed for the data records collected at higher ATP concentration. Variance was calculated for rolling window of 20 ms, at a sampling rate of 10 kHz. Positive position-feedback was used to increase the amplitude of thermal fluctuations, which effectively increased the variance ratio between binding events and free dumbbell noise for both traps in the direction of the actin filament axis. For the examples shown here a combined trap stiffness of 0.07 pN/nm and 0.073 pN/nm and a myosin head stiffness of 2.4 pN/nm and 0.44 pN/nm was estimated for traces in **(A)** and **(B),** respectively. **C)** Scatter plot with mean and standard error of mean (SEM) shows the stiffness measured for ^Sol^M-II and at various ATP concentrations. Each data point displays the stiffness measured from an individual molecule, comprising more than 100 binding events per molecule. **D)** Scatter plot shows stiffness at 1 and 10 μM ATP for ^Pso^M-II. ^Sol^M-II stiffness at 10 μM ATP is presented for comparison. Independent sample t-test was used to calculate the statistical significance. ^Sol^M-II at 10 μM ATP (N = 22) and 500 μM ATP (N = 34) displayed highly significant difference with P < 0.0001. No significant difference was found between ^Sol^M-II and ^Pso^M-II at 10 μM ATP, P = 0.0632. N = number of individual motor molecules.

To further consolidate our observations, we employed another method to estimate the stiffness of ^Sol^M-II as shown in Figure 4A, by applying large triangular wave to both the beads as introduced in Lewallee et al (Lewalle et al., 2008). The acto-myosin binding events were detected during the upward or downward excursion of the dumbbell (Figure 4A). The myosin association with actin restricted the movement of the trap. The force-displacement relations were used to estimate the motor head stiffness (cf. details in methods). Stiffness measured with this alternative method showed similar trend as observed with variance-covariance method, i.e., high (1.51 ± 0.1 pN/nm) and low stiffness (0.55 ± 0.03 pN/nm) at 10 μM and 500 μM ATP, respectively. ^Pso^M-II displayed the stiffness of 1.38 ± 0.13 pN/nm at 10 μM ATP (Figure 4C).

**Figure 4.**
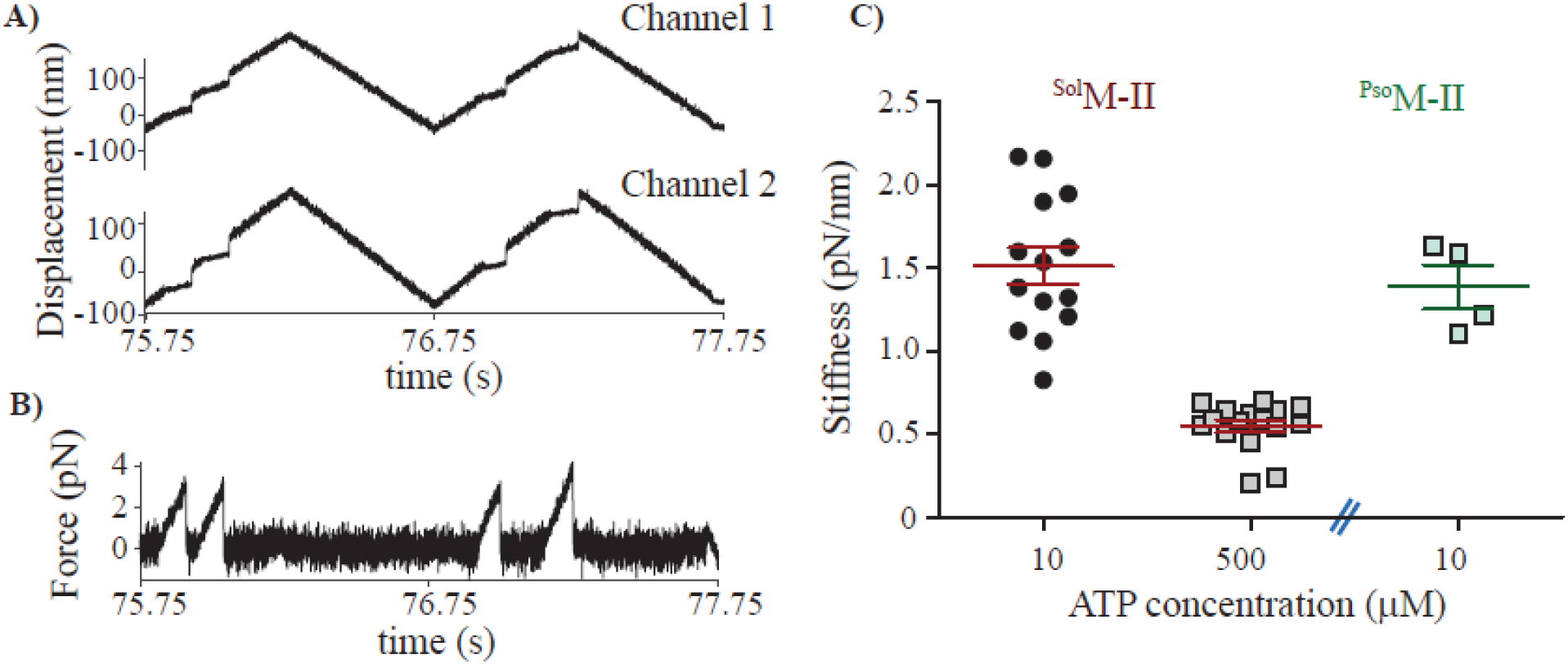
Ramp method used to measure the stiffness of individual motors. **A)** Big triangular wave of 1 Hz with 240 nm amplitude was applied on both the beads (referred as channel 1 and 2). Original data records during interaction of ^Sol^M-II with F-actin at 20 μM ATP. At least 4 binding events observed as steps in the upward ramps and 1 incomplete event toward the end of the downward ramp. **B)** Force experienced by the motor (trace derived from channel 2). **C)** Scatter plot displaying the stiffness of ^Sol^M-II measured at 10 μM ATP (N = 14, n = 960) and 500 μM ATP (N = 16, n = 465) and at 10 μM ATP for ^Pso^M-II (N = 4, n=85). For ^Sol^M-II the mean stiffness at 10 and 500 μM ATP, were significantly different with P < 0.0001 (Independent sample t-test). ^Sol^M-II and ^Pso^M-II, mean stiffness values at 10 μM ATP, are not significantly different, P = 0.30. N = number of individual myosin molecules and n = number of AM association events.

The stiffness values for fast myosin are comparable with previous single molecule measurements with rat fast myosin S1(Capitanio et al., 2006), whereas the high stiffness values for ^Sol^M-II at low ATP concentration reported here are new.

We propose that the two stiffness values represent the two different nucleotide states of the slow myosin head, i.e., AM.ADP state and AM rigor state. The low stiffness corresponding to AM.ADP state is captured at high ATP concentration as the lifetime of the AM rigor state is negligible. Whereas, the rigor stiffness is obtained at low ATP concentration (e.g., 10 μM) as the ADP bound state is very short (on average 30 ms) as compared to the rigor state, comprising ~ 85% of the bound lifetime. Our results suggest comparable rigor stiffness for both fast and slow myosin.

With two acto-myosin cross-bridge states having >2 fold difference in stiffness is it possible to distinguish the change in noise level within the bound period? Detection of such changes would require combination of sufficiently long durations of each state (i.e., state with low and high stiffness). For ^Sol^M-II, we examined several t_on_ events at 10 μM ATP concentration, and found that such changes in noise were indeed detectible. Two such examples are shown in Figure 5. We calculated the stiffness for each noise level, i.e., decrease in noise following the initial binding, yielding two times lower stiffness than that estimated for the duration towards the end of the bound period.

**Figure 5.**
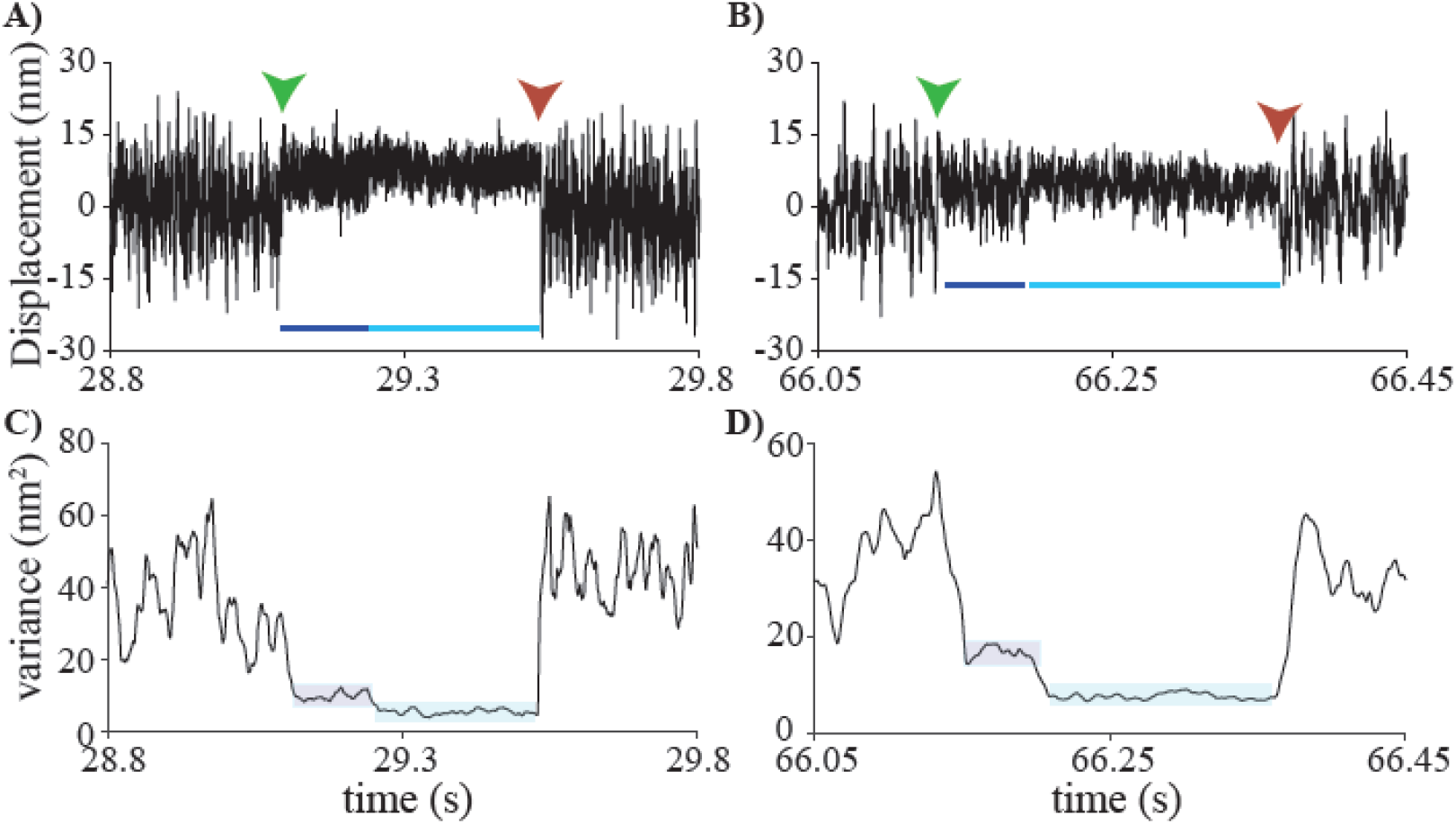
Two noise levels within AM bound state. (**A and B**) Original data records displaying 2 individual AM bound lifetime events of ^Sol^M-II measured at 10 μM ATP. The binding and release of myosin from actin is marked with green and red arrow, respectively. Within the bound duration, time segments with distinct noise amplitudes are indicated with dark and light blue lines, displaying the transition from low to high stiffness of the motor head. **(C and D)** Variance of the corresponding traces in A and B, indicating two levels of variance within an individual binding event, emphasized with light background.

### Single headed myosins -kinetics and mechanics

We further expanded our investigation to understand if the observed differences were specific to the double headed myosins. We generated single headed myosins by papain digestion of full length slow and fast myosins, referred as subgragement-1 (S1), ^Sol^S1 and ^Pso^S1, respectively. Similar to full length myosin, the ^Sol^S1 and ^Pso^S1 displayed a large difference in the actin filament gliding speed (Figure S2). As described in previous section, we measured the duration of acto-myosin bound state, the average displacement of actin by myosin, and the stiffness of the myosin motor at low (10 μM) and high (500 μM) ATP concentrations for ^sol^S1. The average acto-myosin lifetimes were about 5-fold different at the 10 and 500 μM ATP concentrations, i.e., 0.268 ±0.0122 s and 0.053 ±0.001 s as shown in Figure 6 A and B. In agreement with our findings from native slow myosins, the amplitude of stroke size was 6.13 ± 0.15 nm and 4.13 ±0.18 nm at 10 and 500 μM ATP, respectively (Figure 6 D and E). This result further supported that the overall stroke size consists of two distinct sub-steps (4.13 nm and 2 nm), associated with the Pi and ADP release. Yet again, we observed a significant difference in stiffness of the single motor heads during association with actin, i.e., higher stiffness (1.69 ± 0.15 pN/nm) at low ATP and lower stiffness (0.67 ± 0.07 pN/nm) at high ATP concentration using variance-covariance as well as ramp method (i.e., 1.21± 0.15 and 0.43± 0.06 pN/nm) as shown in Figure 6 G and H. Similar to our observation with the full length ^Sol^M-II, we found two distinct noise levels, corresponding to the ADP bound and AM states within the bound durations for ^Sol^S1(cf. Figure S3).

**Figure 6.**
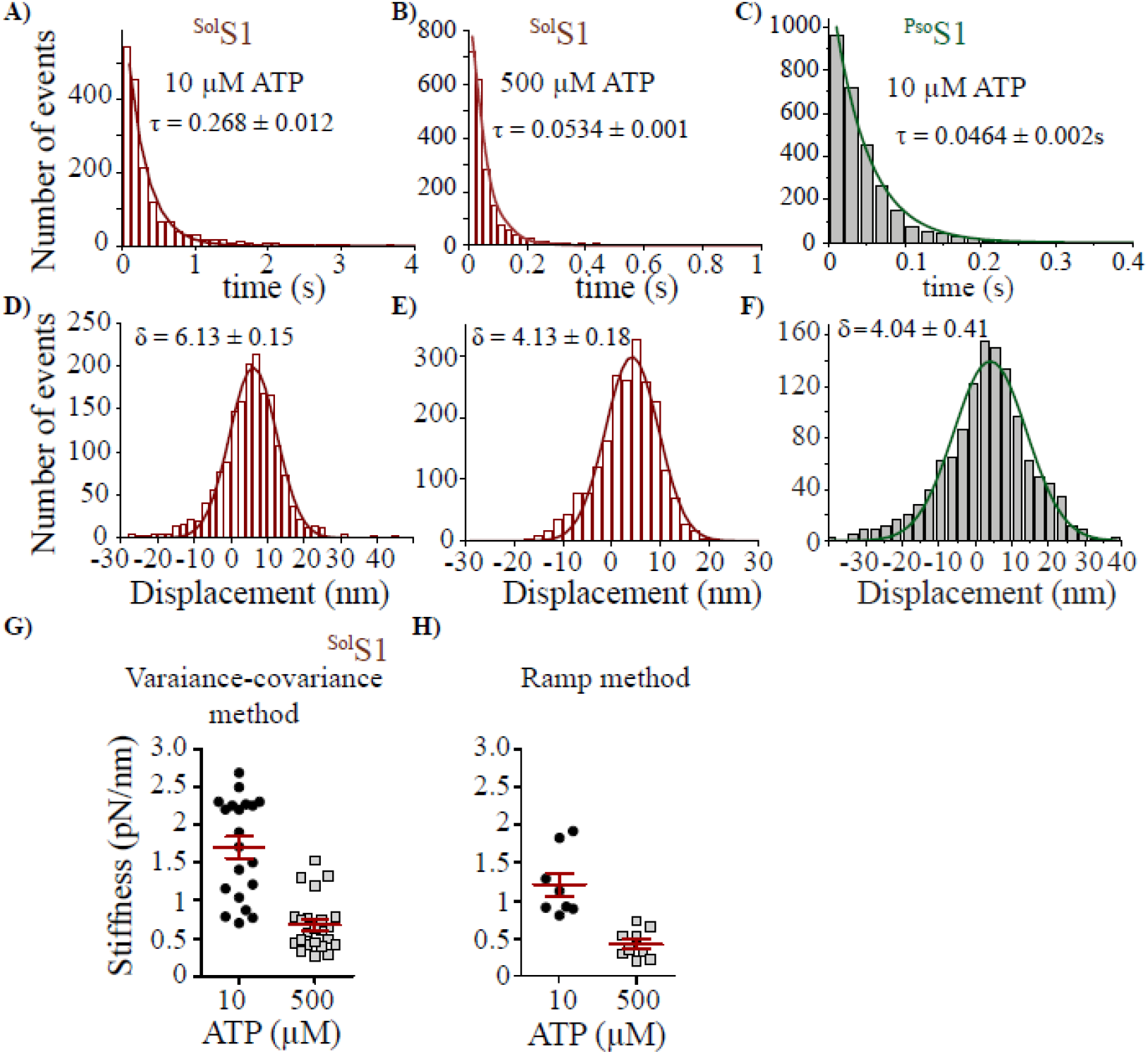
Kinetic and mechanical parameters of ^Sol^S1 and ^Pso^S1. **A - C)** Histograms comprising lifetimes of AM bound states measured at 10 μM and 500 μM ATP concentration for solS1and at 10 μM ATP for ^Pso^S1. The average lifetimes (τ) calculated by least square fitting each histogram with single exponential decay function. **D-F)** The average stroke size (δ) was calculated by least square fitting each histogram with Gaussian function. ^Sol^S1 at 10 μM ATP (N =20, n = 2793), ^Sol^S1 at 500 μM ATP (N=23, n = 1972). ^Pso^S1 (N =20, n =2680) at 10 μM ATP. **G and H)** Stiffness of ^Sol^S1 measured using variance-covariance and ramp method compared. Each data point represents stiffness calculated from individual molecule. Variance-covariance method – same data records as described above for determining the τ and δ, were analyzed, therefore the number of molecules are the same. Ramp method - ^Sol^S1 (N = 8, n=300) at 10 μM ATP, at 500 μM ATP (N = 9, 190). For ^Sol^S1 independent samples t-test yields statistically significant difference between 10 and 500 μM ATP, both for stiffness and stroke size, P < 0.0001. N = number of individual myosin molecules and n = number of AM association events.

For ^Pso^S1, we measured the average acto-myosin lifetime of 0.046 s, the average stroke size of 4.04 nm (Figure 6 C and F) and stiffness of 1.12 ± 0.03 pN/nm at 10 μM ATP. Thus, ^PsoS1^displayed a nearly 1.7- fold slower detachment rate (1/t_on_) of 22 s^−1^, however the mechanical properties remained unaltered in comparison to ^Pso^M-II. These results consolidated our findings with native full length myosins, further indicating that that the cross-bridge stiffnesses arise from single head interaction with actin.

Altogether, the difference in kinetic and mechanical properties displayed by ^Pso^M-II and ^Sol^M-II, were recapitulated in their single headed motors.

## Discussion

The objective of this study was to directly measure the kinetic and mechanical properties of native dimeric single myosin molecules that are responsible for observed differences in the performance of fast *vs.* slow muscle. Under defined ATP concentrations, the individual acto-myosin interactions were probed to measure the detachment rates, stroke size and cross-bridge stiffness. A major finding from this study was the similar acto-myosin crossbridge rigor stiffness for ^Pso^M-II and ^Sol^M-II. Importantly, for ^Sol^M-II, we could demonstrate that the nucleotide state of myosin determines the measured stiffness. The ‘AM.ADP’ state with stronger actin affinity, displayed ~2-fold lower stiffness as compared to the ‘rigor’ state, the state with strongest actin affinity. The overall powerstroke occurring in two substeps was also detectible by adjusting the ATP concentrations, such that the duration of strong bound ‘ADP’ or the ‘rigor’ states dominated the duration of the measured AM bound states. Thus, apart from the overall stroke size, the 1^st^ powerstroke associated with Pi release, and even the conformational change linked with ADP release (2^nd^ powerstroke) was identified for ^Sol^M-II. Due to weak ADP affinity of the ^Pso^M-II, the 2^nd^ powerstroke associated with the ADP release could not be resolved, meaning rather complete strokesize was measured. Other observed differences included > 10- fold faster AM detachment rates and shorter stroke size for the fast ^Pso^M-II in comparison to ^Sol^M-II.

Studies on chicken skeletal and smooth muscle myosin reported higher force and displacement for the dimeric myosins than the single headed motor forms (Tyska et al., 1999). Important to note that lower ATP concentrations under their experimental conditions may have supported simultaneous double head interaction with actin. On the contrary, we found that the mechanical properties (stroke size and stiffness) of the single headed myosin (S1) were similar to the dimeric full length M-II for both examined isoforms. Therefore, our observation further reinstates the previous idea that the single myosin head engages in the AM cross-bridge cycling even for the dimeric myosin II. The kinetic properties, i.e., the AM detachment rate however differed by almost 60%, i.e., the dimeric form dissociated faster compared to the single headed myosin, measured at specific ATP concentration. This finding further suggests that although a single head associates with the actin filament, the second motor head influences the AM detachment rates, presumably by facilitating optimal AM interactions.

In summary, both fast and slow myosin forms possess a similar cross-bridge stiffness in rigor states. Therefore, our results support that primarily the AM detachment rate determines actin filament velocity, which is limited by either the ADP release rate for the ^Sol^M, while for ^Pso^M the rate of ATP binding determines the AM detachment kinetics under our experimental conditions and thereby the gliding velocity.

Our measured cross-bridge stiffness for ^Pso^M-II is in close agreement with previous studies performed at single molecule and single fiber level (Capitanio et al., 2006), (Lewalle et al., 2008),(Brenner et al., 2012, Percario et al., 2018, Linari et al., 2007). In previous studies, the measured stiffness ranged from 0.6 - 1.7 pN/nm for the fast myosin alone, using different experimental approaches (cf. Table S1). For ^Sol^M-II, however, the high cross-bridge stiffness of ~1.5 pN/nm was never reported. Interestingly, lower values of measured stiffness (~0.5 pN/nm) were reported earlier for ^Sol^M-II from different species, rabbit, mouse, and human, both in single-molecule and fiber studies as ‘rigor’ stiffness (Capitanio et al., 2006, Percario et al., 2018, Brenner et al., 2012). We assigned the lower value of cross-bridge stiffness to the AM.ADP state. Our results imply that the transition from ‘ADP bound’ to ‘rigor’ state leads to the structural rearrangement of the lever arm relative to the catalytic domain resulting in approximately 2 nm actin displacement that accompanies the change in the elasticity of the cross-bridge (schematized in Figure 7). In fact, the two distinct rigidities of the same isoform of motors were never reported to our knowledge.

**Figure 7.**
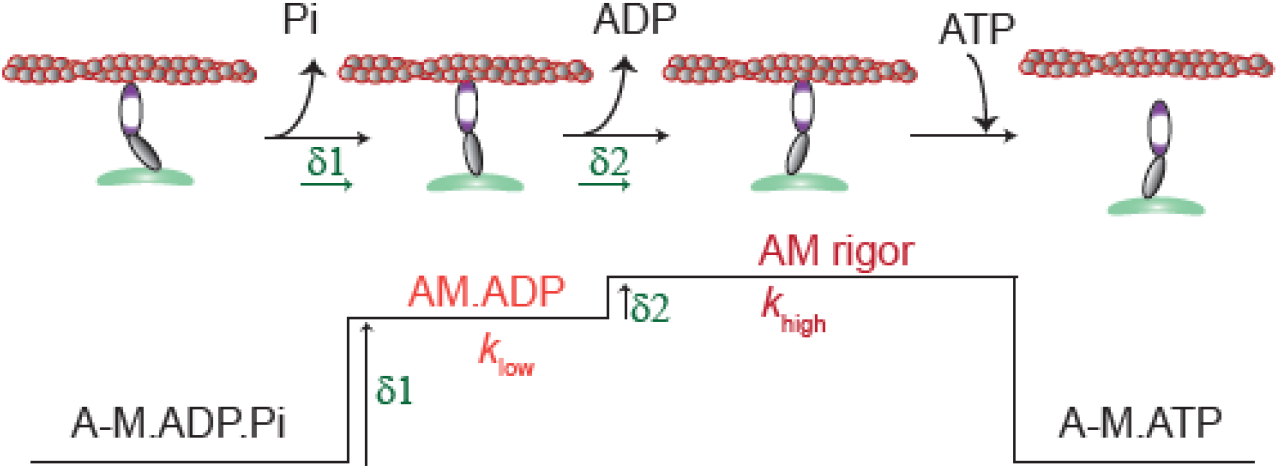
Mechanism of chemo-mechanical coupling. During acto-myosin cross-bridge cycle - the release of ATP hydrolysis products Pi and ADP from myosin active site leads to the conformational changes in the motor head causing swinging of the lever arm in 2 sub-steps, larger followed by relatively smaller stroke size associated with Pi and ADP release, respectively. The AM.ADP state with lower myosin head stiffness as compared to the AM rigor state is identified for the slow myosin. *k*_*low*_- low stiffness, *k*_*high*_- high stiffness, *δ1* and *δ2* - 1^st^ and 2^nd^ stroke, A-actin, M - myosin, Pi - inorganic phosphate. Weakly bound or detached state is indicated with a hyphen between A and M. Strongly bound AM states are indicated with red and dark red labels.

For several myosin isoforms, the ADP release step was previously shown to be strain sensitive (Veigel et al., 2003, Veigel et al., 2005). It is plausible that the lower stiffness of the AM.ADP state is a key mechanical feature for the strain sensing mechanism of the myosin, and furthermore essential for the structural change triggering ADP release and the second powerstroke. Although, this feature was not identified in other myosins until now, it would be opportune to examine the motor forms that displayed strain sensitivity.

Apart from the slow and fast muscle type-specific mechanical differences, intriguingly, under certain experimental conditions, Brenner and colleagues observed that the mechanical parameters barely differed between fast and slow fibers. The stiffness-speed relations measured in rigor conditions were essentially the same for human soleus fibers compared to rabbit psoas fibers.

Only at the slowest speeds of stretch, a decrease in rigor stiffness for slow fibers was detectible (Kohler et al., 2002, Brenner, 1991, Brenner et al., 2012). Besides, for soleus fibers, they found a temperature-dependent change in active fiber stiffness and cross-bridge strain. Stiffness during isometric contraction measured at a lower temperature (5°C) was two-fold lower as compared to the measurements at 20°C. In contrast, for psoas fibers, such temperature-dependent change was negligible (Brenner et al., 2012). The authors attributed the observed change to increased occupancy of force-generating states.

Recent X-ray diffraction and EM studies highlighted the different arrangements of myosin molecules in the thick filaments in rat slow and fast fibers, i.e., the simple lattice *vs.* super lattice configuration, respectively (Ma et al., 2019). This study further indicated that super lattice configuration in the fast fibers aids the accessibility of the actin filament for myosin molecules as compared to the simple lattice arrangement. This implied that fast fibers could readily engage the motors and thereby generate higher force than slower fibers.

Thus, these studies and our results provide clues that under defined experimental conditions, factors such as the cross-bridge densities, different force-generating states, or even change in force produced per head need to be considered when estimating the single myosin molecule stiffness in different muscle fibers.

We observed high variability of the stiffness values measured from ^Sol^M-II myosin molecules (0.6 −2.8 pN/nm), with an average stiffness of ~1.5 pN/nm. A possible reason may be the different immobilization of the myosin molecules on the nitrocellulose surface, both for full length and S1. To ensure that pure population of slow ^Sol^M-II are employed for stiffness estimation, as an internal control, we checked the lifetimes of the ‘t_on_’ states for the corresponding myosin molecules. Please note that the event duration ‘t_on_’ for ^Sol^M-II was ~7 times longer than that of fast myosin at the 10 μM ATP concentration. At ATP concentration of more than 20 μM, the AM interaction events for the fast myosin were too short to resolve with our set-up. The soleus muscle primarily contains (about 97%) slow myosin II isoform. Fast myosin isoform comprising ~3 % as a contaminant could be excluded using this approach. In our ensemble molecule gliding assays, 3% contaminant motors are unlikely to influence the actin filament speed. Nearly 20% contaminant motors would be required to cause a substantial change in the velocities, as previously concluded from experiments with a mixture of different ratios of myosin isoforms (Cuda et al., 1997, Harris and Warshaw, 1993).

We used full length dimeric motors, where both myosin heads may form cross-bridge under low ATP concentrations, meaning the high stiffness could arise from simultaneous binding of 2 heads of myosin to the adjacent actin monomers. Such interactions would result in sequential binding and release of the two heads. However, following systematic analysis of the AM bound periods in data records, we did not observe two step unbinding at the end of the bound periods. Moreover, our stiffness and stroke size measurements using single headed myosin - S1 under same experimental condition strengthened the conclusion that only one of the two heads associated with actin and thus contributed to the measured stiffness. Our observations are in agreement with previous suggestions drawn from muscle fiber studies, which reasoned in favor of a single myosin head interaction with actin during cross-bridge cycle (Offer and Elliott, 1978, Huxley and Tideswell, 1997).

Although the main elastic region of the myosin that contributes to the cross-bridge stiffness remains unknown, the probable regions include the lever arm (light chain binding domain)(Uyeda et al., 1996, Dobbie et al., 1998, Irving et al., 2000) and the converter region that links the catalytic domain to the lever arm. Brenner and colleagues showed a ~3 fold increase in the cross-bridge stiffness of the beta cardiac myosin heavy chain (β-MHC) due to single point mutations in the converter region (Arg719Trp). Therefore, the converter region was proposed to be the main elastic element in the motor head (Kohler et al., 2002, Seebohm et al., 2009). It is, however, difficult to endorse the identity of an elastic element in the myosin head from our study.

The fast and slow myosin isoforms share nearly 80 % amino acid sequence homology in the head domain. Even for the different species, such as mouse, rabbit, pigs, and humans sequence difference of nearly 20% is found between the converter region of slow and fast myosins. The differences in various domains between the two isoforms are discussed in more detail in Pellegrino et. al. (Pellegrino et al., 2003). Our results raise other interesting possibilities to reflect on the lower measured forces in previous measurements i.e., amino acid differences in the myosin head influencing the actin binding properties or the ATPase activity, such that the longer duration of the AM.ADP transition state (i.e., a state with low cross-bridge stiffness in our measurements) in slow myosin, corresponds to the lower force generation during the AM cross-bridge cycle. The extrapolation to muscle fiber studies, however, requires careful consideration as several differences in single molecule *v*s. motor stiffness in single fiber measurements exist. Further independent experiments in fibers will be necessary to address the effects observed at single molecule level.

## Conclusion and outlook

In summary, our studies revealed previously unexplored aspect of the myosins mechanical properties. Accordingly, the transition from AM.ADP to AM rigor state generates a 2^nd^ powerstroke of defined size and is associated with ~two-fold change of stiffness of the myosin head. Knowledge of the elasticity of the motors has deeper implications to understand the molecular mechanism of the force generation. Functional diversity among myosin isoforms or among various motor types due the ATPase kinetics influencing the mechanics or vice versa is key to gain comprehensive understanding of the molecular basis of the motor function and their diverse physiological role. The information is also vital to comprehend certain myopathies related to mutations in specific domains of the myosin. Besides, the switch in myosin isoform expression under pathological condition or during development is expected to affect the chemo-mechanical features. Furthermore, the precise insights into the motor properties could be relevant for the manipulation of the biochemical/biomechanical states as a target for the therapeutic intervention.

## Material and methods

### Native myosin II and generation of myosin Subfragment-1(S1)

Full length rabbit fast skeletal muscle myosin II, ^Pso^M-II, and slow myosin, ^Sol^M-II was isolated from skinned fibres of *M. psoas* and *M. soleus* muscle in an extraction buffer (0.5 M NaCl, 50 mM HEPES, pH, 7.0, 5 mM MgCl2, 2.5 mM MgATP, and 1 mM DTT) as previously described (Amrute-Nayak et al., 2008, Thedinga et al., 1999). Isolated myosin was aliquoted, flash frozen in liquid nitrogen and stored at −80°C in 50% glycerol. To generate single headed subfragment-1(myosin S1), both ^Pso^M-II and SolM-II myosin isoforms were digested with papain (myosin S1), as described earlier (Margossian and Lowey, 1982, Trybus and Chatman, 1993). S1 was stored at −80°C in 50% glycerol in a buffer (5 mM Na-phosphate, 10 mM K-acetate, 4 mM Mg-acetate and, 2 mM DTT at pH 7.0). To collect the muscle tissues, the New Zealand white rabbits, Crl:KBL (NZW) were euthanized as per the guidelines from German animal protection act §7 (Tötung zu wissenschaftlichen Zwecken/ sacrifice for scientific purposes). In this study, we used shared organs originating from the animals approved for experiments with authorization number 18A255. The animal registered under reference number G43290, was obtained from Charles River France. All the procedures were carried out in accordance with relevant guidelines and regulations from the Lower Saxony State Office for Consumer Protection and Food Safety and Hannover Medical School, Germany.

### Preparation of actin filaments

For in vitro actin filament motility assays, actin filaments were generated by incubating chicken G-actin in polymerisation buffer (p-buffer) containing 5 mM Na-phosphate, 10 mM K-acetate, and 4 mM Mg-acetate, supplemented with protease inhibitor overnight at 4°C. Equimolar concentration of fluorescent phalloidin was added to fluorescently mark the actin filaments. Unlabelled actin filaments were prepared without any phalloidin. To get long biotinylated actin filaments (≥ 20 μm) for optical trapping experiments, G actin and biotinylated G actin was mixed in equimolar ratios to final concentration of 0.1 μg/ μl each in p-buffer containing 1 mM DTT, 1 mM ATP, and 0.5 mM protease inhibitor, AEBSF (Cat No 30827-99-7 (PanReac Applichem ITW). The mixture was incubated overnight at 4°C, and followed by addition of fluorescent phalloidin to label the actin filaments.

### Active myosin heads

Prior to use in *in vitro motility* assays or optical trapping measurements, the myosin S1 was further purified to discard any inactive motors by co-sedimentation with concentrated F-actin. Actin-myosin complex was dissociated by addition of 2 mM MgATP. While the active heads released from actin in the presence of ATP, the inactive myosin remained bound and separated by centrifugation at 70,000 rpm for 30 min at 4°C. The supernatant containing the enzymatically active motors was further supplemented with 2 mM DTT and protease inhibitor and used in the functional assays. This procedure was routinely followed prior to the main experiments.

### In vitro motility assay

*In vitro motility* assay was performed with full length fast ^Pso^M-II and slow ^Sol^M-II by immobilisation of the motors on the bovine serum albumin (BSA) coated surface, while monomeric ^Pso^S1 or ^Sol^S1 was immobilised on nitrocellulose coated surface, respectively. The assay is described in more details in (Amrute-Nayak et al., 2008). Briefly, myosin was incubated for 5 min on BSA or nitrocellulose coated surface, followed by wash and surface blocking with 1 mg/ml BSA in assay buffer (AB; 25 mM imidazole hydrochloride pH- 7.2, 25 mM NaCl, 4 mM MgCl2, 1 mM EGTA, and 2 mM DTT). To block the inactive or damaged myosin motor heads 0.25 μM short, unlabelled F-actin was injected in the flow cell and incubated for 1 min. 2 mM ATP was introduced in the chamber to release the actin filaments and to make the active motor heads accessible. ATP was washed out with AB buffer. TMR (tetra methyl rhodamine) labelled F-actin was incubated for 1 min, followed by a washing step to remove excess filaments. Finally, the chamber was infused with buffer containing 2 mM MgATP and antibleach system (18 μg/ml catalase, 0.1 mg/ml glucose oxidase, 10 mg/ml D-glucose, and 10 mM DTT) to initiate F-actin motility. Images were acquired with a time resolution of 200 ms (i.e., 5 frames/sec) using a custom-made objective-type TIRF microscope. Actin filament gliding speed was analysed with Manual Tracking plug-in from ImageJ (NIH, Bethesda, USA).

### 3-bead assay with optical tweezers

The optical trapping set up is described previously (Steffen et al., 2001, Steffen and Sleep, 2004). For the assay, flow cells with approximately 15 μl chamber volumes were assembled using coverslips with nitrocellulose coated beads. Glass microspheres (1-1.5 μm) suspended in 0.1 % nitrocellulose in amyl acetate were applied to 18×18 mm coverslips. All the dilutions of biotin-actin filaments and myosin S1 were made in reaction buffer containing 25 mM KCl, 25 mM Hepes (pH 7.4), 4 mM MgCl2, and 10 mM DTT. The full length native myosin was diluted in high salt extraction buffer without MgATP. For the experiment, the chamber was prepared as follows, 1) flow cells were first incubated with 1 μg/ml native myosin or myosin S1 for 1 min, 2) followed by wash with 1 mg/ml BSA and incubated further for 1 min to block the surface, 3) finally, reaction mixture containing 0.8-1μm neutravidin coated polystyrene beads and 1-2 nM biotinylated actin was flowed in with 10 μMATP (or varied concentrations of ATP), ATP regenerating system (0.01 mM creatine phosphate, and 0.01 unit creating kinase) and deoxygenating system (1 mg/ml catalase, 0.8 mg/ml glucose oxidase, 2 mg/ml glucose, and 20 mM DTT). The assembled flow chamber was sealed with silicon and placed on an inverted microscope for imaging and trapping.

An actin filament was trapped in between the two laser trapped beads (Figure 1A), pre-stretched, and brought in contact with the 3^rd^ bead immobilized on the chamber surface. Low-compliance links between the trapped beads and the filament were adjusted such that the ratio of the position variance during free and bound periods was in the range of 4-10 as described in Smith et al.(Smith et al., 2001). The bead positions were precisely detected with two 4-quadrant photodetectors (QD), recorded and analyzed. The acto-myosin interaction events were monitored as a reduction in free Brownian noise of the two trapped beads. Data traces were collected at a sampling rate of 10,000 Hz and low-pass filtered at 5000 Hz.All the experiments were carried out at room temperature of approximately 22°C. For the fast myosin - to improve the time resolution and detect short-lived AM binding events, high-frequency triangular wave of ~600 Hz and about 120 nm amplitude was applied to one of the trapped beads as described in (Amrute-Nayak et al., 2019) (Veigel et al., 2002).

### Data Analysis

Using the variance-Hidden-Markov-method (Smith et al., 2001), acto-myosin interactions events detected as reduction in noise were analyzed. This method allowed the AM bound states (low variance) to be distinguished from unbound states (high variance). Matlab routines were employed to evaluate data records for ‘AM’ interaction lifetime, ‘*t_on_*’, stroke size (*δ*) of motors.

Two different methods were employed to measure the stiffness (*k*) of the myosin motors, namely - variance-covariance method and ramp method as described in detail previously (Smith et al., 2001) (Lewalle et al., 2008). To calculate motor stiffness with the variance-covariance method, data records with combined trap stiffness in the range 0.05–0.09 pN/nm were used.

Briefly, for the ramp method, a big triangular wave was applied on both the beads. A large amplitude triangular wave of 240 nm and 1 Hz was chosen to study acto-myosin binding events at low ATP concentration (10 μM) due to their long life times, while 120 nm amplitude and 2 Hz wave was applied at high ATP concentrations of 500 μM ATP. Thus, constant ramp velocity of ~500 nm/s was used for high and low ATP concentrations.The AM interaction events were registered on both upwards or downwards sides of the ramp. The beads follow the movement of the trap in myosin unbound state, whereas the binding event restricts the bead movement by exerting restraining force, thereby reducing the velocity of bead movements. For the AM binding events, the velocity ratios between unbound and bound states for left and right beads were calculated. The AM cross-bridge stiffness was deduced using the velocity ratios, trap stiffness and combined link stiffness as described previously in Lewalle et al.(Lewalle et al., 2008).

### Single myosin molecule interaction with actin filaments in optical trapping measurements

To ensure that each data record is derived from an intermittent interaction between a single myosin head and actin, myosin density on the bead surface was adjusted by diluting the myosin solution. We standardised a routine in our measurements whereby, typically, one bead was found to interact with the dumbbell among 8-10 beads scanned for the presence of motor on the bead. This routine minimized the likelihood of multiple molecules simultaneously interacting with the actin filaments. From the Poisson distribution, in our measurements we estimate the likelihood of presence of more than 1 motor per bead to be less than 1 %. From a total of 259 beads we analyzed for AM binding events; we do not expect that a few beads with more than 1 head could alter our results.

### Statistical analysis

All values are expressed as mean ± SEM and indicated in the manuscript in relevant sections. Poisson distribution was used to estimate the likelihood of more than 1 molecule interacting with actin filaments in optical trap measurements. Gliding velocities, single motor stroke size, and stiffness of the native and S1 were analyzed using independent samples t-test. The nonparametric Mann-Whitney U test was used to calculate the statistical differences in the duration of AM interaction events, *t*_on_ for the fast and slow motors. Statistical significance was assumed if P < 0.05.

## General

We thank Petra Uta for the technical assistance in protein preparations, Prof. Theresia Kraft, Dr. Ante Radocaj, Dr. Tim Scholz, and Dr. Walter Steffen for critical comments on the manuscript.

## Author contributions

MA conceived the project and designed the experiments with inputs from TW, BB and AN. TW performed optical trapping measurements and analyzed the data. AN performed protein preparations. AN and MA performed motility experiments and data analysis. Motility and single molecule data was interpreted by TW, AN, and MA. MA wrote and edited the manuscript with assistance from TW and AN.

## Data and materials availability

Please contact MA for any material requests.

## Supplementary Information

**Figure S1.**
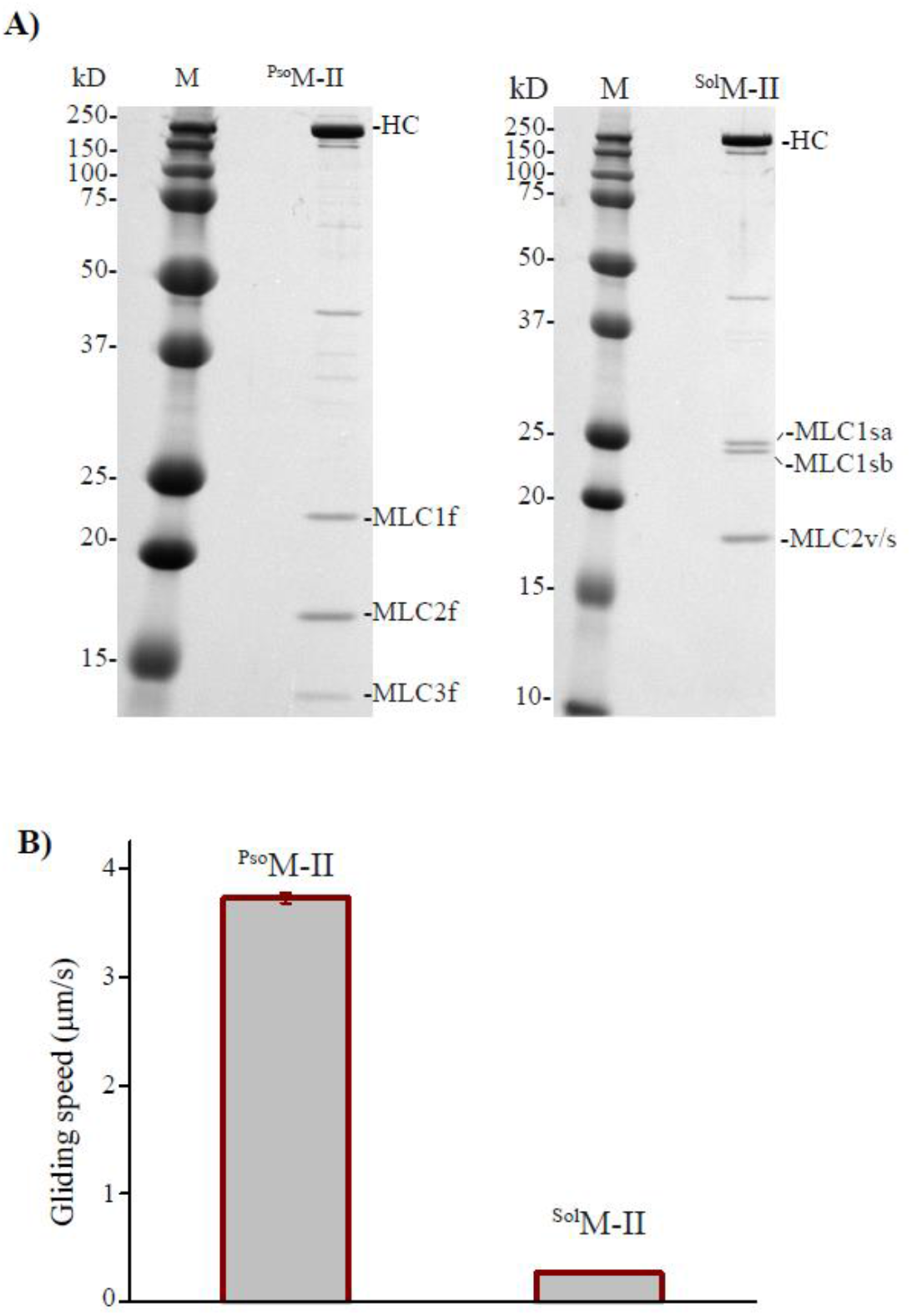
**A)** Gel images display native full length myosins extracted from rabbit M. psoas and M. soleus muscle fibers. ^Pso^M-II lane displays the 2-native ELC isoforms, a long (~21 kDa, MLC1f) and a short one (~16 kDa, MLC3f) and RLC (MLC2f). ^Sol^M-II lane also display 2-distinct ELC isoforms (MLC1sa and MLC1 sb with molecular weight of 27 and 24 kDa, respectively), and a RLC isoform, MLC2v/s. The myosin probes were resolved on 12.5 % SDS-PAGE gel and stained with Coomassie blue stain. M-protein marker in the gel images. **B)***In vitro actin filament gliding.* The gliding speed of ^Pso^M-II and ^Sol^M-II is compared in a bar diagram. N ≥ 100 filaments each. Error bars – SEM. Please note that the error bars for ^Sol^M-II is not visible as the value is too small. All motility experiments were repeated minimum three times with independent myosin preparations with similar results. \**P* < 0.0001; independent samples *t*-test.

**Figure S2.**
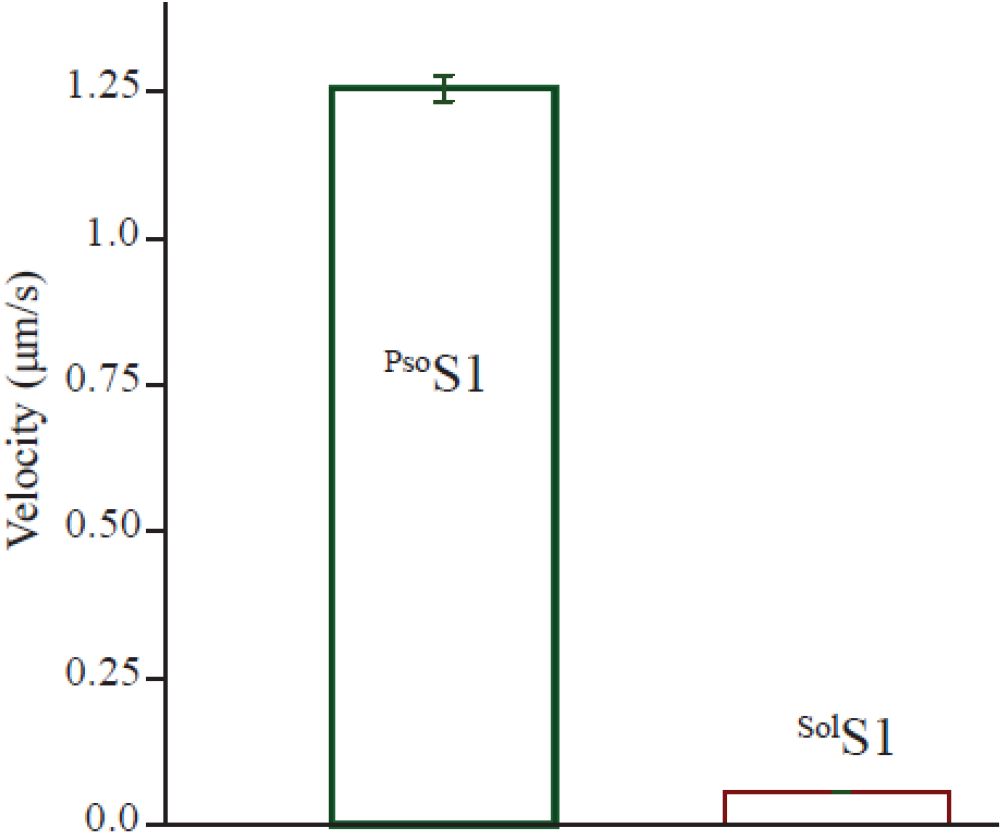
In vitro actin filament gliding. Gliding speed of ^Pso^S1and ^Sol^S1 is compared in a bar diagram. The experiments were performed at 2 mM ATP and 22°C. Minimum 3 different myosin S1 preparations were tested with highly reproducible motility results. N ≥ 100 filaments each analyzed. *P* < 0.0001; independent samples *t*-test. Error bars – SEM

**Figure S3.**
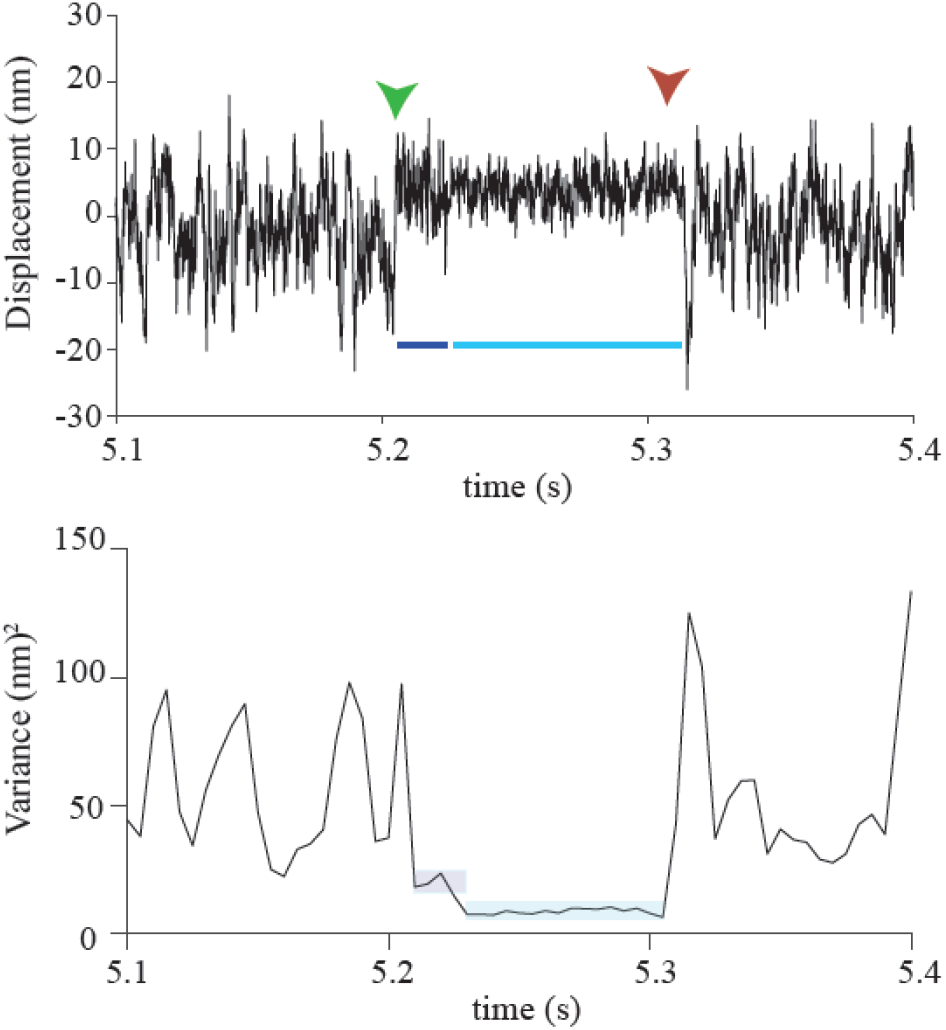
Single actomyosin interaction trace with distinct noise levels. Upper panel shows original data records measured at 10 μM ATP displaying an individual AM interaction event of SolS1, as evident by the reduction of noise. The binding and release of myosin from actin is marked with green and red arrow, respectively. Time segments within the bound duration displayed distinct noise amplitudes, marked with dark and light blue lines, which indicate the transition from low to high stiffness of the motor head. Lower panel shows variance of the corresponding trace. Two levels of variance within an individual binding event are emphasized with light background.

**Table S1.**
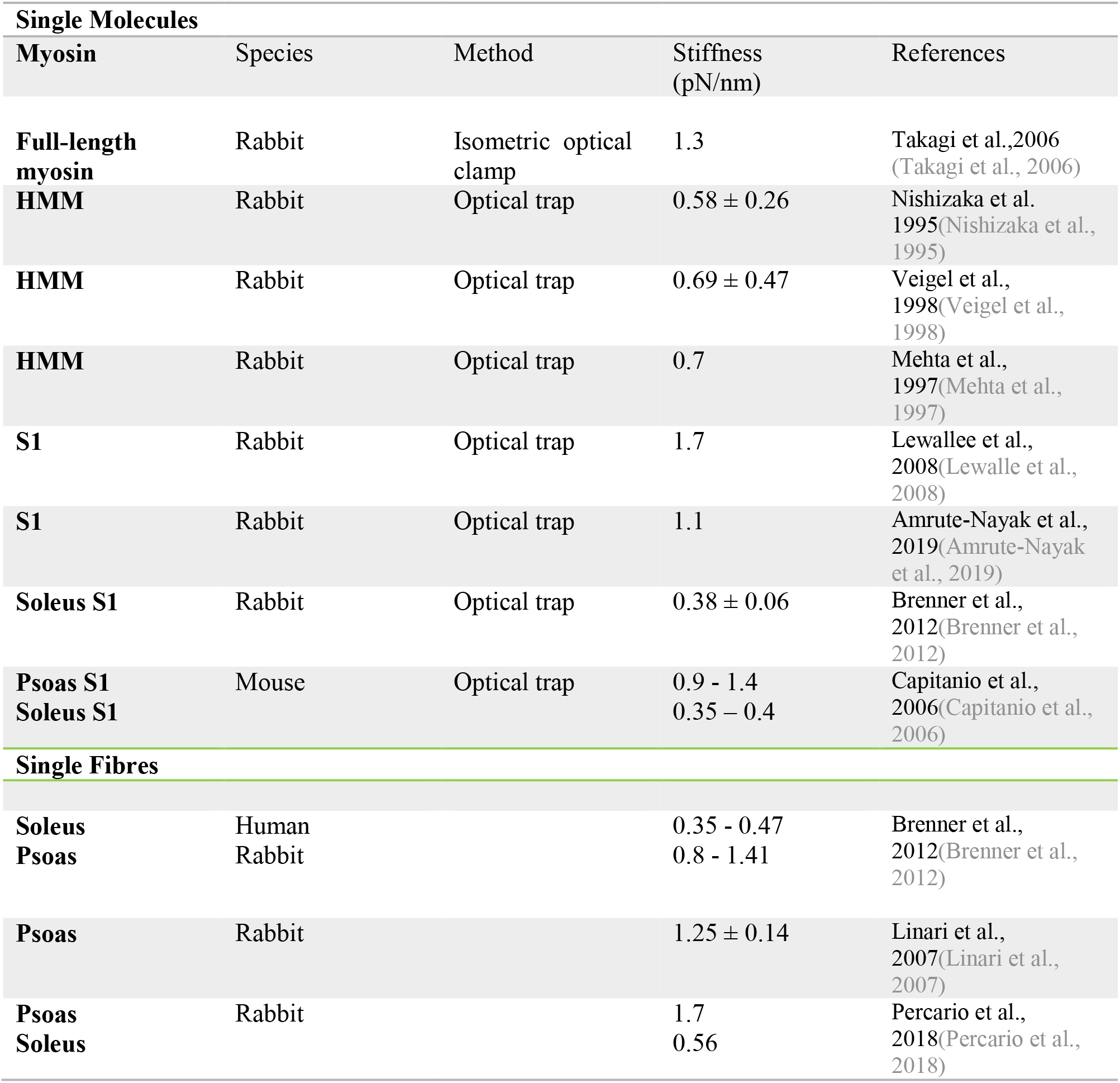
Stiffness of myosin molecules from previous studies is summarised. Single myosin head stiffness estimated from either direct measurement of single myosin motors or estimated from fibre studies. Full length myosin – dimeric native myosin, HMM – dimeric heavy meromyosin, S1-single headed subfragement-1.

